# LCP1 Regulates Cell Motility in Chondrosarcoma and Correlates with Metastatic Potential and Poor Patient Outcomes

**DOI:** 10.1101/2023.01.31.526513

**Authors:** Caleb Watson, John T. Martin, Makoto Nakagawa, Nicholas Guardino, Trudy Zou, Emily Peairs, Alexandra Krez, Puvi Nadesan, ZeYu Huang, Jianhong Ou, Jason A. Somarelli, Benjamin Alman, Julia D. Visgauss

## Abstract

Chondrosarcoma (CSA) is the second most common primary malignancy of bone, whose aggressive potential and chemo-resistant nature result in extremely poor outcomes in patients with advanced disease. Grading and prognostication of these tumors remain a significant challenge for pathologists, and medical oncologists have no effective therapies to prevent or treat metastatic disease. In the present study, we sought to explore the pathogenesis and aggressive progression of chondrosarcoma by comparing gene expression differences from metastasizing and non-metastasizing CSA tumors. We hypothesized that metastasizing tumors have inherent differences that may be attributed to the dysregulation of specific oncogenic cell signaling pathways. RNA-Seq analysis from patient-derived low-passage cell lines of metastasizing and non-metastasizing tumors pinpointed the gene *LCP1* (lymphocyte cytosolic protein 1) as upregulated in CSA cells from metastasizing tumors. Analysis of a large clinical cohort of CSA demonstrated that increased expression of *LCP1* correlated with poor patient survival. *In vitro* analyses confirmed the ability of LCP1 to promote migration and invasion of CSA cells. These data support the key role of LCP1 on metastatic potential and poor prognosis in CSA. In conclusion, we confirm the ability of LCP1 to drive metastatic behavior and correlate with poor outcomes in patients.

## Introduction

Chondrosarcomas (CSA) are malignant tumors classified by their production of cartilage matrix[1]. They are the second most common primary malignancy of bone, with an incidence of about 1 in 200,000 every year[2]. Central conventional CSA accounts for the vast majority of subtypes, comprising 85-90% of all CSA[1]. Currently, histological grading is the main predictor of clinical behavior and patient prognosis. Grade is determined by evaluation of the degree of cellularity, nuclear atypia, and mitoses[1, 3]. Classification consists of grade 1 (low-grade) CSA, which has a ten-year survival rate of 83%, grade 2 (intermediate-grade) CSA, with a ten-year survival rate of 64%, and grade 3 (high-grade) CSA, with a ten-year survival rate of 29%[1]. De-differentiated CSA are high-grade tumors with undifferentiated histologic appearance and lack of cartilage matrix production that arise within a central conventional CSA of a lower grade. Five-year survival for this advanced subtype is estimated at only 18%[4].

While histological grading remains the standard for predicting metastatic risk in CSA, it is highly subject to inter-observer variability, with debate in the literature and amongst pathologists regarding grading metrics. As a result, grading is poorly predictive of the metastatic potential of any given tumor. Standard treatment for CSA is wide, *en-bloc* resection[1]. Unfortunately, there exists no effective systemic therapies to prevent or treat metastatic disease, and the prognosis in this setting is dismal. Miao et al. (2019) reported a median overall survival of 13.9 months in their cohort of 72 patients with de-differentiated CSA, with distant metastasis common during or after initial presentation[5].

Given the inherent challenges with both the classification and treatment of malignant CSA, there is an urgent need to identify and better understand the molecular mechanisms associated with metastatic progression in CSA to improve prognostication and identify novel targeted systemic therapies.

Tumor metastasis often involves a process by which malignant cells escape the primary tumor, migrate to secondary sites, and adapt to the distant tissue microenvironment, acquiring necessary traits to avoid extracellular death signals and promote further macroscopic growth[6]. Prior research in CSA has failed to identify prognostic biomarkers or genetic drivers of disease, and we believe that the pathogenesis and aggressive progression of CSA is driven by dysfunction in post-transcriptional regulation of complex molecular pathways. We hypothesize that metastasizing and non-metastasizing CSA have inherent differences that may be attributed to the dysregulation of specific oncogenic drivers and cell signaling pathways.

To investigate this process, we established a panel of low passage primary cell cultures from the primary tumors of patients with CSA, including patients for whom the disease metastasized and for whom the disease did not metastasize. Analysis of these cell lines by bulk RNA-Seq identified high expression of lymphocyte cytosolic protein 1 (*LCP1*) in tumors that metastasized. *In vitro* studies confirmed a direct effect of LCP1 on migratory and invasive ability of CSA cells. Furthermore, evaluation of RNA-Seq data from a large patient chondrosarcoma cohort confirmed the prognostic relevance of *LCP1* expression in patients with CSA, with increased levels of *LCP1* correlating with decreased survival.

## Methods

### Cell lines

In this study we used deidentified patient-derived primary cell cultures from our institution’s biobank. These were derived from the primary tumors of patients with histologically confirmed intermediate and high-grade CSA. Clinical information about each patient tumor included the metastatic status of the patient at diagnosis and most recent follow-up, confirmed with at least 5-years follow up. Consent from patients was obtained at the time of tumor resection for bio-banking with clinical data and use of tissue in preclinical studies per institutional IRB approved protocols. A total of five different cell cultures were used throughout this study. The following three cell lines were from patients who developed metastatic disease: 711, 725, 731. The following two cell lines were from patients who remained disease free: 732, 743. While all of these cell lines were generated from primary tumors, we refer to these cell lines as “metastasizing” and “non-metastasizing” respectively, based on the metastatic status of the patient from which they were derived.

### RNA-Sequencing

RNA-seq analysis was conducted by the Duke Sequencing and Genomic Technologies Core to determine differentially expressed genes (DEGs) between metastasizing and non-metastasizing cell lines (met: 711, 725; non-met: 743, 732). To ensure quality control, RNA-seq reads were trimmed by Trim Galore (version 0.4.1, with -q = 15). These reads were then mapped using the TopHat algorithm (version 2.1.1, with parameters -b2-very-sensitive-no-coverage-search and supplying the UCSC hg38 known Gene annotation), which aligned reads on the basis of the human genome (GRCh38) without having to rely on known splice sites[7]. The mapped reads were filtered by a map quality scores (MAPQ) no smaller than 30 which were then counted using featureCounts (version 1.6.4)[8]. The Bioconductor package DESeq2 (version 1.28.1) was employed to analyze differential expressions (DE) in the mapped reads[9]. Coverage depth was normalized by deeptools (version 3.1.3) using the Reads Per Kilobase per Million mapped reads (RPKM) for RNA-seq (Ramirez et al. 2016). Transcripts Per Million (TPM) values were quantified from Salmon (version 1.2.1) quantification and summarized through tximport (version 1.16.1)[10, 11]. Of note, summary statistics included in supplemental tables reflect FeatureCount read quantification.

Volcano plots were created using ggplot, with dysregulation defined by ± 2 log2-fold change (WithShrink) and p-value < 0.05[12]. Text was added using geom_text_repel methods within R, labeling those distinct points with ± 5 log2-fold change (WithShrink) and p-value < 1E-10. Heatmaps were created using the heatmap.2 function within the ggplots package. In these plots, calculated Z-scores for each row (gene) as well as normalized counts from each well were compared between those annotated genes from the volcano plot in triplicate across each of the four cell lines[13]. Dendrograms were applied for analysis of hierarchical clustering of cell line replicates and individual genes.

### qPCR

Quantitative PCR (qPCR) was performed to further validate genes identified from the RNA-seq in four cell lines: two metastasizing (711, 725) and two non-metastasizing (732, 743). RNA concentration and quality were quantified using the NanoDrop 2000/2000c program. 1050 ng of RNA was isolated from each cell line and combined with RNAse free water to a volume of 16.8 μL. Finally, 4.2 μL iScript 5x was added for a final reaction volume of 21 μL. The cycling conditions were priming for 5 minutes at 25°C, reverse transcription for 20 minutes at 46°C, and reverse transcription inactivation for one minute at 95°C. qPCR reactions were performed in triplicate in a 384-w plate using Quantstudio 6Flex System for amplification. Reaction mixtures were composed of the following: 1.6 μL cDNA, 0.5 μL each of forward and reverse primer, 2.4 μL RNAse free H2O, and 5.0 μL 2x SsoAdvanced Universal SYBR Green Supermix (BioRad) for a reaction mixture for 10.0 μL.

Primers for the genes of interest were custom designed using the IDT PrimerQuest Tool, with optimum amplicon size of 100 bp and optimum melting temperature of 60°C. The primer sequences used in qPCR are listed here:

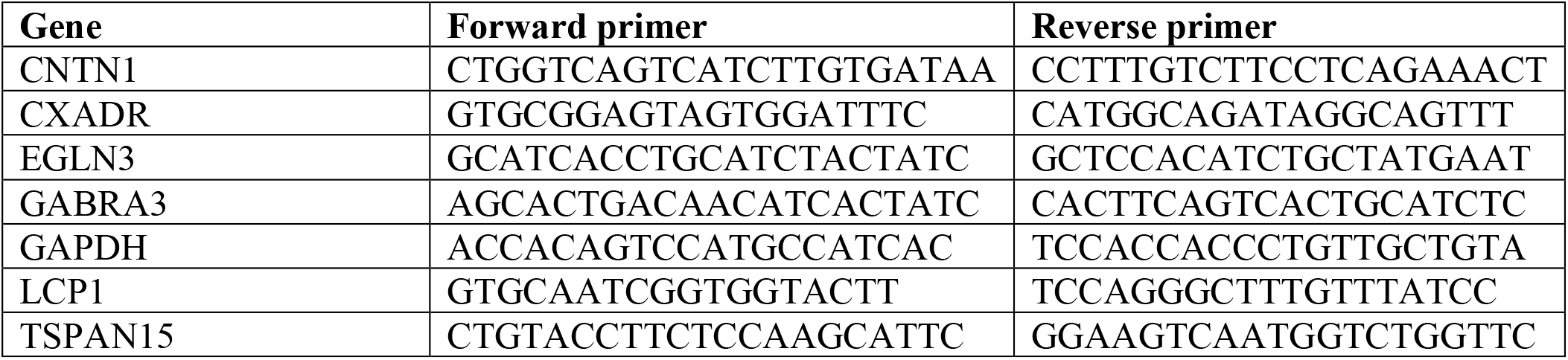

Fold change was calculated by normalization to GAPDH internal control (IDT ReadyMade™ forward and reverse primers), and significance of expression was assessed using one-way ANOVA and post-hoc student’s t-test with p-adj < 0.001.

### Western blotting

Proteins were extracted from cultured cells using RIPA Lysis and Extraction Buffer (Pierce) with Halt Protease and Phosphatase Inhibitor Cocktail (Pierce). The samples were centrifuged at 18,800 *g* for 10 min at 4 °C. The BCA colorimetric method was performed to check the concentration of the proteins. Cell lysates were electrophoresed using Mini-PROTEAN TGX Stain-Free Protein gels (Biorad) and transferred onto Immune-Blot PVDF membranes (Biorad). After blocking nonspecific binding sites with 5% nonfat milk in 0.1% Tween 20/Tris-buffered saline (TBS-T) for 1h, the membranes were treated at 4 °C overnight with the following primary antibodies in blocking buffer: anti-LCP1 (MA5-11921; Thermo Fisher); anti-Vinculin (MAB3574; Millipore); anti-GAPDH (60004-1-Ig; Proteintech); and anti-HA High Affinity (11867423001; Roche). After washing, the membranes were incubated with horseradish peroxidase-conjugated secondary antibodies (1:1,000, 7074, 7076, 7077, Cell Signaling) for 1 hour at 4 °C. Immune complexes were visualized by Clarity Western ECL Substrate (Biorad) or SuperSignal West Femto Maximum Sensitivity Substrate (Thermo Scientific). Images were analyzed by Image Lab software (Biorad).

### LCP1 overexpression

The pLV-EF1a-*LCP1*-HA-IRES-Puro construct was generated by inserting cDNA encoding HA-tagged *LCP1* into the lentiviral vector pLV-EF1a-IRES-Puro. pLV-EF1a-IRES-Puro was a gift from Tobias Meyer (Addgene plasmid # 85132; http://n2t.net/addgene:85132; RRID: Addgene_85132)[14]. Ecotropic lentivirus was produced using Lenti-X 293T cells (Takara). 743 and 732 cells were transfected with pLV-EF1a -*LCP1*-HA-IRES-Puro construct together with psPAX2 and pMD2.G using GeneJuice (Millipore), and the supernatants containing lentivirus were collected after 48 h. Both psPAX2 (Addgene plasmid #12260; http://n2t.net/addgene:12260; RRID: Addgene_12260) and pMD2.G (Addgene plasmid #12259; http://n2t.net/addgene:12259; RRID: Addgene_12259) were gifts from Dr. Didier Trono. 743 and 732 cells were then infected with lentiviruses for 24 h using Polybrene (final concentration of 10 μg/mL; Sigma) in high-glucose DMEM supplemented with 10% FBS and 1% P/S. Infected cells were selected with puromycin for 48 h prior to cell viability assay and wound healing assay.

### LCP1 Knockout with CRISPR/Cas9

The lentiCRISPRv2 vector expressing three Single-guide RNA (sgRNA) targeting *LCP1* gene was generated by Duke Core Facility. LentiCRISPRv2 was a gift from Feng Zhang (Addgene plasmid # 52961; http://n2t.net/addgene:52961; RRID: Addgene_52961)[15]. Ecotropic lentivirus was produced using Lenti-X 293T cells (Takara). These cells were transfected with lentiCRISPRv2 vector expressing sgRNA together with psPAX2 and pMD2.G using GeneJuice (Millipore), and the supernatants containing lentivirus were collected after 48 h. 725 and 731 cells were then infected with lentiviruses for 24 h using Polybrene (final concentration of 10 μg/mL; Sigma) in high-glucose DMEM supplemented with 10% FBS and 1% P/S. Infected cells were selected with puromycin for 48 h and genetic deletion was validated by western blot analysis against LCP1. Two LCP1-knockout cell lines were cultured in high-glucose DMEM supplemented with 10% FBS and 1% P/S and subjected to cell viability assay and wound healing assay.

### Cell viability assay

The effect of LCP1 knockout and overexpression on cell viability was evaluated using CellTiter-Glo 2.0 Cell Viability Assay (Promega). Briefly, 5.0 × 10^3^ cells were seeded in each well of a 96-well plate in 100 μl high-glucose DMEM supplemented with 10% FBS and 1% P/S (day 0) and allowed to adhere overnight. At 24 h after seeding, the cell viability was analyzed for 5 days in all the cells using Spark (TECAN).

### Wound healing assay

Cell motility was assessed by a wound healing assay. Briefly, 5.0 × 10^4^ cells were seeded in each well of a 96-well plate in 100 μl high-glucose DMEM supplemented with 10% FBS and 1% P/S and allowed to grow overnight to form a spatially uniform monolayer. After creating the scratch by WoundMaker (Essen BioScience), the wells are washed twice with culture media and 100 μl fresh medium was added to each well. The plate was placed into the IncuCyte Live-Cell Analysis System (Essen BioScience) and allowed to equilibrate for 5 min. Images of each well were scanned every two hours for a total duration of 24 h (725, 743, 732 cells) or 48 h (731 cells). Wound Width was calculated with IncuCyte software to evaluate cell motility.

### ORIS invasion assay

Cell migration was interrogated using an Oris 96-well cell migration assay kit (Platypus Technologies, Madison, WI) per manufacturer instructions. Briefly, cells were plated at a concentration of 3 x 10^5 cells per well. After removing stoppers, cells were allowed to migrate into the empty hydrogel field for 72 hours. Images of wells were taken every 8 hours at 4x using an Incucyte S3 (Satorius, Göttingen, Germany) until the completion of the experiment. Images were captured using fluorescence filters to identify endogenously labeled cells (GFP or mCherry). The total number of cells which migrated into the empty hydrogel field were counted using ImageJ. Statistical testing was performed in GraphPad Prism v9. For two-group comparisons, unpaired t-tests were used; p<0.05 conferred statistical significance for all tests.

### Unsupervised learning of CSA gene expression phenotypes in a clinical cohort

A total of 84 malignant CSA with mRNA expression and overall survival data in a publicly available database[16] were reviewed. This database was authorized by the Advisory Committee on Information Processing in Material Research in the Field of Health (CCTIRS) and the French Data Protection Authority (CNIL). CSAs from the above dataset were clustered on mRNA expression data and patient survival was subsequently evaluated by cluster. After pre-processing microarray data using an existing protocol in R (R v4.0.2; Bioconductor [v3.13]), dimensionality reduction was performed using principal components analysis (prcomp, stats package [v3.6.2]; n=60 components [90% cumulative variance]) and was followed by hierarchical clustering (agnes, cluster package [v2.1.2]; Ward’s method, Euclidean distance, n=3 groups). Kaplan-Meier survival curves (survfit, survival package [v3.2-11]) and hazard ratios (coxph, survival package) were then generated for each cluster.

To examine the clinical impact of the expression level of these upregulated DEGs, a median-split was performed for each differentially expressed gene, followed by Kaplan-Meier survival curves and hazard ratios for tumors with high and low expression levels. Survival analysis was further analyzed using quartile level of expression for LCP1.

## Results

### RNA-Seq reveals differential expression of LCP1 between metastasizing and non-metastasizing tumors

We performed RNA-Seq on chondrosarcoma primary cell cultures derived from the primary tumors of patients who developed metastatic disease (725, 711) and patients who did not develop metastatic disease (743, 732). This analysis identified 1421 genes with statistically significant differences in the CSA cell lines where the patient developed metastasis (905 upregulated and 516 downregulated) (Figure 1A; Supplemental Tables 1 and 2). Gene expression patterns clustered the samples derived from metastatic and non-metastatic patients (Figure 1B).

**Figure 1.**
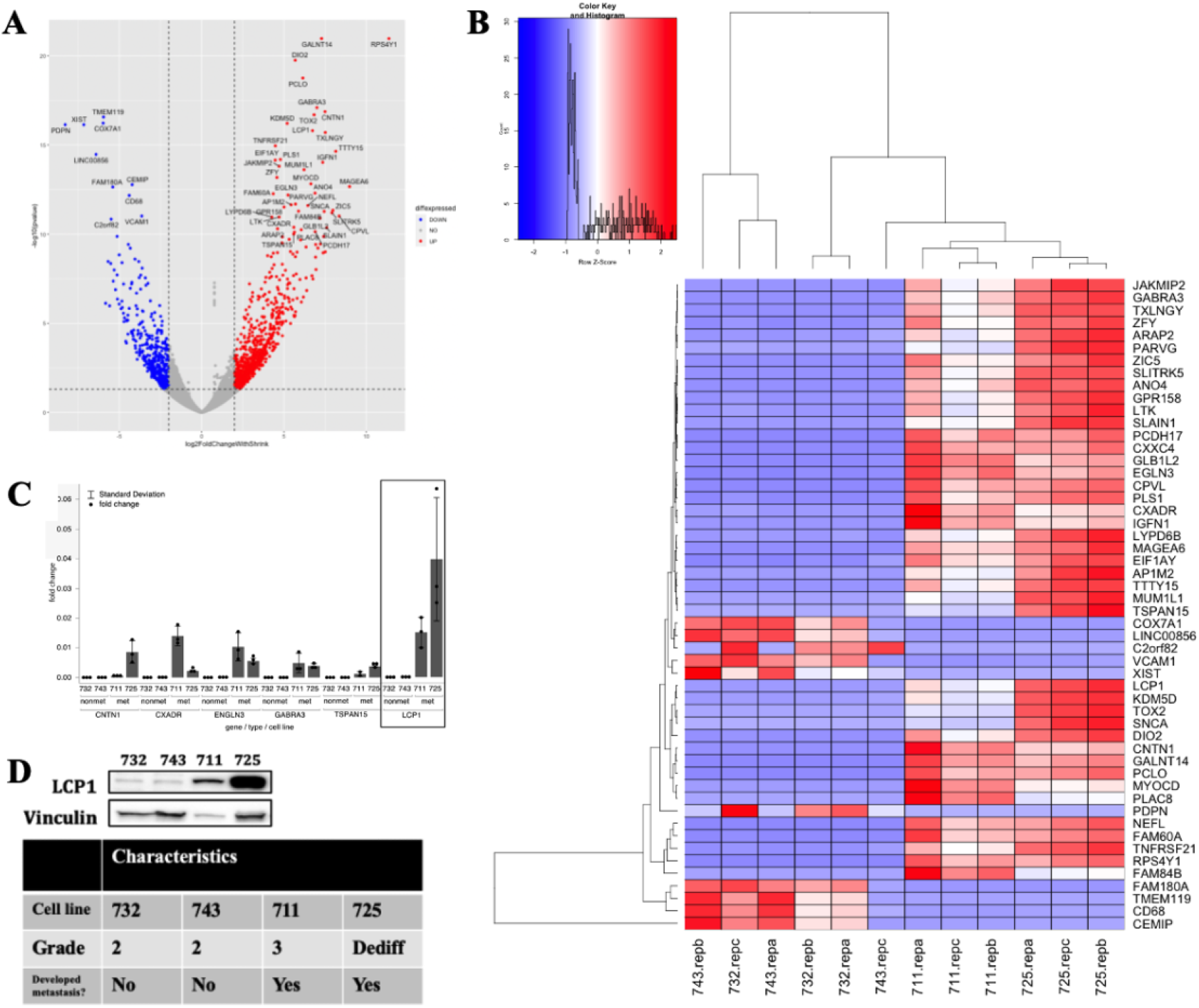
Correlation of LCP1-upregulation to metastatic potential in CSA. (A) Volcano plot showing dysregulated genes between CSA cells from primary tumors of patients with metastatic disease (711, 725) and those who remained disease free (732, 743) at final follow up. Labelled thresholds set at p-value < 0.05 and ± 2-log2FC (with shrink) and annotated thresholds set at ± 5-log2FC (with shrink) and p-value < 1E-10. (B) Heatmap comparing raw counts in triplicate (clustering by column). Genes listed constitute those DEGs labelled on the volcano plot. (C) qPCR validation of gene expression, showing individual data points for each cell line replicate, performed on selected cohort of significantly upregulated genes from RNA sequencing. LCP1 stands out as most differentially expressed in CSA cells from patients who developed metastatic disease. (D) Western blot confirming increased protein levels of LCP1 in human CSA cells from patients with metastatic disease.

We further validated by qRT-PCR a subset of the top differentially-upregulated mRNAs from the RNA-Seq results, including *CNTN1, CXADR, EGLN3, GABRA3, LCP1, and TSPAN15*. All genes were confirmed to exhibit upregulation in the chondrosarcoma cells from the patients who developed metastasis, with the greatest difference observed for *LCP1* (Lymphocyte Cytosolic Protein 1) (Figure 1C). Western blotting again confirmed high protein levels of LCP1 in CSA tumor cells from patients who developed metastasis, and low LCP1 protein expression in CSA tumor cells from patients who did not develop metastasis (Figure 1D).

### LCP1 mediates cell motility of CSA cells in vitro

The substantial upregulation of *LCP1* prompted us to further investigate this gene in chondrosarcoma cells *in vitro*. Given the previously established role of LCP1 in lymphocyte activation and migration, we hypothesized that LCP1 would affect migration and invasive ability of chondrosarcoma cells (Morley et al. 2012). To do this, we first used CRISPR/Cas9 to knockout *LCP1* in the CSA cells with high endogenous levels of LCP1: 725 and 731 (Figure 2A). Consistent with our hypothesis, *LCP1* knockout cells showed prolonged wound closure compared to wild type cells (725: p = 0.003, 731: p < 0.001; Figure 2C). Additionally, LCP1 knockout cells exhibited decreased invasive potential (decreased cell numbers in the detection zone) as compared to wild type cells in Oris invasion assays (725: p < 0.001, 731: p < 0.01; Figure 2D-E). We further confirmed the effect of LCP1 on cell motility by ectopically expressing an HA-tagged *LCP1* in the CSA cells with low endogenous levels of LCP1: 732 and 743 (Figure 3A). *LCP1* overexpressed cells demonstrated faster wound closure (732: p < 0.001, 743: p = 0.002; Figure 3C), and increased invasive potential (increased cell numbers in the detection zone) compared to wild type cells in Oris invasion assays (732: p < 0.001, 743: p < 0.001; Figure 3D-E). These results were not due to differences in proliferation, as no significant difference was observed in cell proliferation/viability between the *LCP1* wildtype cells and their genetically-modified conditions *in vitro* (Figures 2B and 3B).

**Figure 2:**
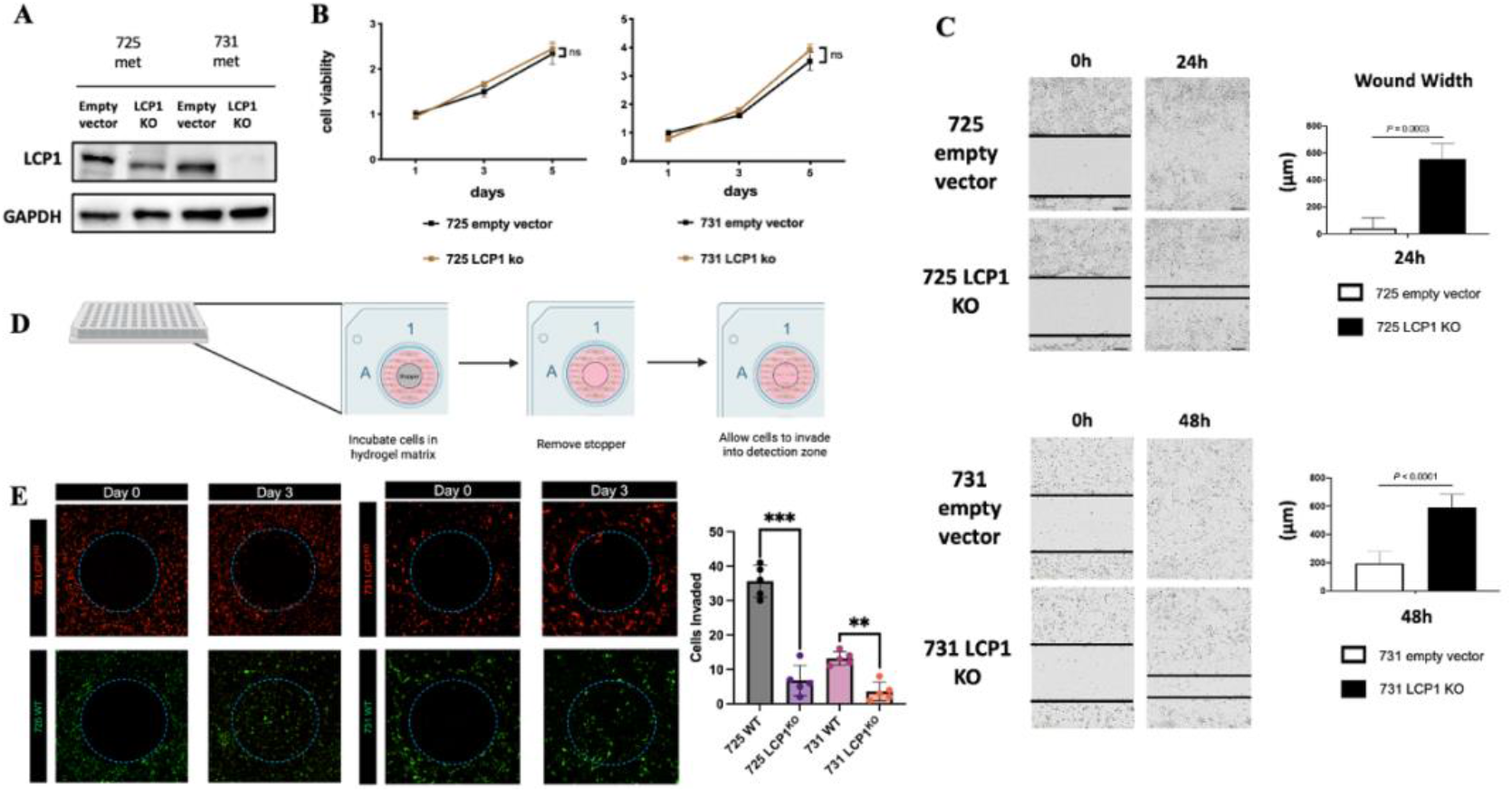
*LCP1* knockout decreases chondrosarcoma cell motility *in vitro*. (A) CRISPR based knockout of *LCP1* revealing decreased protein levels of LCP1. Β Cell viability remains unchanged despite *LCP1* knockout in 725 and 731 cell lines. (C) Scratch wound assay demonstrating decreased motility and delayed wound closure of CRISPR mediated *LCP1* knockout cells compared to wild type in endogenously high LCP1 expressing chondrosarcoma cells. (D) Schematic of ORIS workflow. (E) ORIS cell invasion assays reveal decreased invasive potential after *LCP1* knockout compared to wild type of endogenously high expressing chondrosarcoma cells.

**Figure 3:**
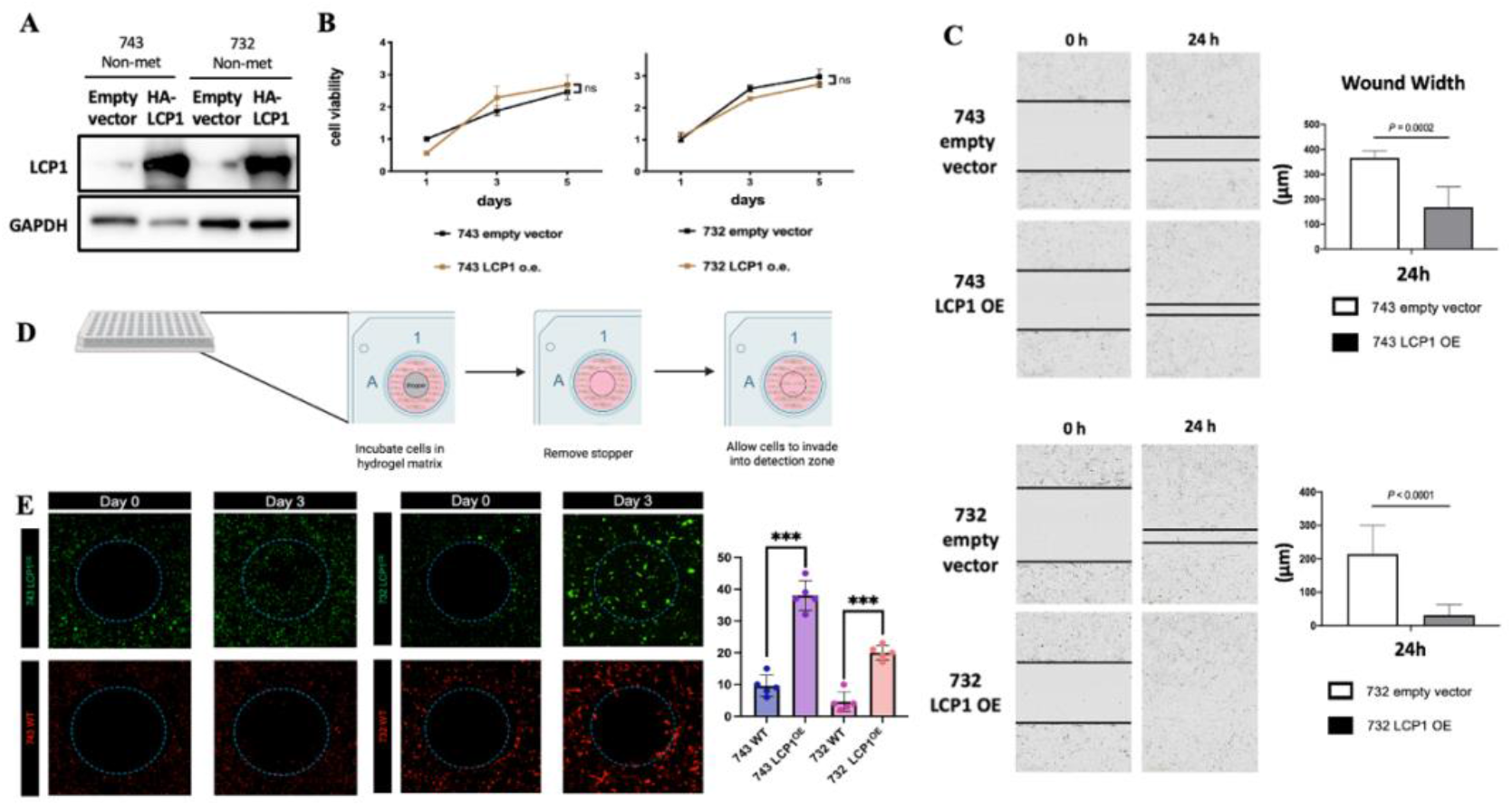
*LCP1* overexpression facilitates aggressive behavior of chondrosarcoma cells *in vitro*. (A) Overexpression of *LCP1* revealing increased protein levels of LCP1. Β Cell viability remains unchanged despite *LCP1* overexpression in 743 and 732 cell lines. (C) Scratch wound assay demonstrating increased motility and wound closure of *LCP1* overexpressed cells compared to wild type endogenously low expressing LCP1 chondrosarcoma cells. (D) Schematic of ORIS workflow. (E) ORIS cell invasion assays reveal increased invasive potential after *LCP1* overexpression compared to endogenously low expressing chondrosarcoma cells.

### Elevated LCP1 expression in CSA tumors correlate with poor survival outcomes in patients

We next investigated the impact of *LCP1* expression on CSA clinical outcomes. Using data from a large publicly-available patient cohort (Nicolle at el. 2019), we performed an unbiased clustering analysis to identify groups of tumors with similar gene expression patterns. Clustering analysis of the CSA tumors within this cohort revealed three distinct clusters with different histological and survival phenotypes. Survival analysis of the clusters revealed distinctly poor survival of cluster 3 tumors (HR vs. C1 = 18.8, p<0.01) (Figure 4A). Cluster 1 and 2 tumors were composed primarily of low to intermediate grade tumors (G1/G2/G3/dediff.; cluster 1 27%/45%;/18%/11%; cluster 2 9%/55%/27%/9%), while cluster 3 tumors were primarily high-grade, dedifferentiated tumors (G1/G2/G3/dediff.; cluster 3 0%/1%;/1%/80%) (Figure 4B). The highest expression of *LCP1* is exhibited within the cluster 3 tumors (Fig. 4C).

**Figure 4.**
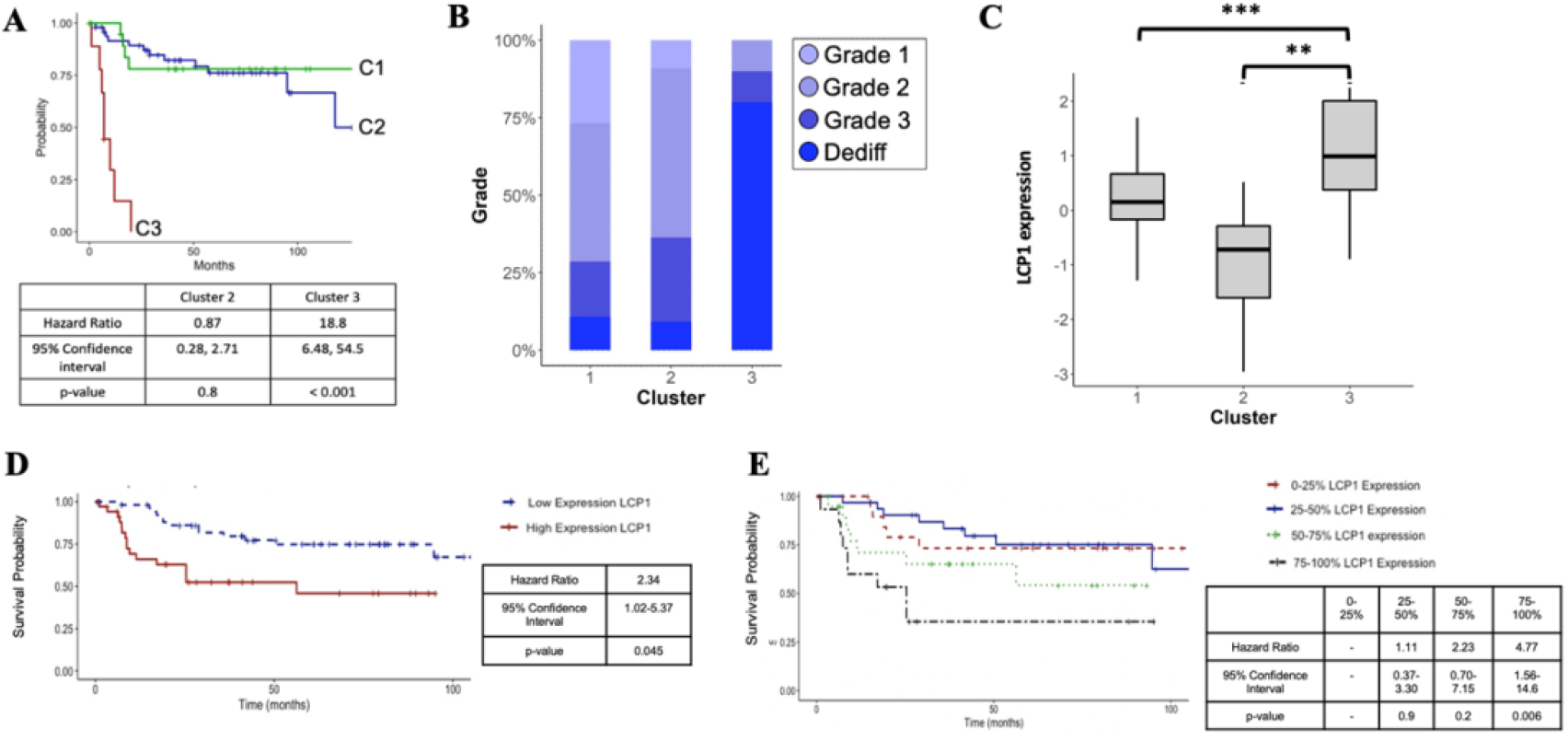
Clustering analysis of chondrosarcomas using mRNA expression from a large patient cohort reveals a tumor subset with poor survival outcomes. (A) Survival outcomes by cluster. (B) Histological grade by cluster. (C) Cluster 3 chondrosarcoma tumors identified with poor survival also have increased expression of *LCP1*. P-value between C1 and C2 < 1E-3, C2 and C3 < 1E-4. Survival analysis using a clinical cohort (Nicolle, et al) reveals increased tumor expression of *LCP1* correlates with poor survival in patients with chondrosarcoma, (D) using median split to define high and low mRNA expression (E) mRNA expression stratified by quartile.

Kaplan-Meier survival curves revealed decreased overall survival in patients with *LCP1* expression above the median (Figure 4D). Additional survival analysis revealed an expression level-dependent effect on survival when stratified by quartile expression levels; *LCP1* expression above the median (50-75, and >75%) revealed incremental decreases in overall survival (Figures 4E).

## Discussion

The inability to overcome intrinsic chemoresistance in CSA has prompted us to focus on better understanding the biology of these tumors in an attempt to identify therapeutic vulnerabilities. This is especially critical in high-grade tumors that are prone to metastasis[17]. There is no current effective treatment for metastasizing CSA, emphasizing the urgent need to 1) understand the specific molecular mechanisms that drive metastatic disease, and 2) identify novel targeted therapeutics. In this paper, we describe our integration of RNA-Sequencing, *in vitro* studies, analysis of clinical data, highlighting LCP1 as an important driver of metastatic potential and disease progression in chondrosarcoma.

LCP1, also known as L-plastin (LPL) and plastin-2, is an intracellular protein critical for actin cross-linking. Peptides derived from LCP1 have been shown to promote changes in cell functions, such as integrin activity and cell adhesion, suggesting that LCP1 may play a role as an adaptor or scaffolding protein critical in both cell motility and cell signaling[18, 19].

The majority of research on LCP1 exists in the context of normal and disordered immune function, as LCP1 function was historically thought to be specific to immune cells, regulating integrin functions, stabilizing immune synapses, and critical for motility and extravasation to target tissues[18, 19]. However, LCP1 has recently been described in several solid organ malignancies, also correlating with aggressive behavior and disease progression. Interestingly, the current body of literature suggests that LCP1 may act both directly through its effects on cytoskeletal organization, and indirectly through activation of downstream oncogenic cell signaling. In prostate cancer, *LCP1* expression is regulated by the PI3K/AKT pathway and positively correlates with lymph node metastases[20]. Upregulation of *LCP1* was found to induce metastasis of osteosarcoma via degradation of NRDP1 and activation of the JAK2/STAT3 pathway[21]. LCP1 has also been found to be exclusively activated in multiple myeloma cells with TRAF3 loss-of-function mutation, the most frequent NF-κB mutation in MM (Shin et al. 2020). Finally, Ser5 phosphorylation of LCP1 was found to be modulated by ERK/MAPK and PI3K activation promoting breast cancer cell invasiveness[22].

Our work has shown that increased expression of *LCP1* in chondrosarcoma correlates with metastatic phenotype and poor patient survival. In addition to providing a novel marker of biologic aggressivity in these tumors, we have also demonstrated that it has direct effects on cell migration and invasiveness, important components of the metastatic cascade. Future work is needed to understand the role of LCP1 more completely on cell behavior and signaling, and the dysregulation that contributes to increased expression in metastasizing tumors.

## Supporting information

Supplemental Table 1- Upregulated

Supplemental Table 2- Downregulated

## References

1. Gelderblom, H., et al., The clinical approach towards chondrosarcoma. Oncologist, 2008. 13(3): p. 320–9.

2. Giuffrida, A.Y., et al., Chondrosarcoma in the United States (1973 to 2003): an analysis of 2890 cases from the SEER database. J Bone Joint Surg Am, 2009. 91(5): p. 1063–72.

3. Evans, H.L., A.G. Ayala, and M.M. Romsdahl, Prognostic factors in chondrosarcoma of bone: a clinicopathologic analysis with emphasis on histologic grading. Cancer, 1977. 40(2): p. 818–31.

4. Strotman, P.K., et al., Dedifferentiated chondrosarcoma: A survival analysis of 159 cases from the SEER database (2001-2011). J Surg Oncol, 2017. 116(2): p. 252–257.

5. Miao, R., et al., Prognostic Factors in Dedifferentiated Chondrosarcoma: A Retrospective Analysis of a Large Series Treated at a Single Institution. Sarcoma, 2019. 2019: p. 9069272.

6. Gupta, G.P. and J. Massague, Cancer metastasis: building a framework. Cell, 2006. 127(4): p. 679–95.

7. Trapnell, C., L. Pachter, and S.L. Salzberg, TopHat: discovering splice junctions with RNA-Seq. Bioinformatics, 2009. 25(9): p. 1105–11.

8. Liao, Y., G.K. Smyth, and W. Shi, featureCounts: an efficient general purpose program for assigning sequence reads to genomic features. Bioinformatics, 2014. 30(7): p. 923–30.

9. Love, M.I., W. Huber, and S. Anders, Moderated estimation of fold change and dispersion for RNA-seq data with DESeq2. Genome Biol, 2014. 15(12): p. 550.

10. Patro, R., et al., Salmon provides fast and bias-aware quantification of transcript expression. Nat Methods, 2017. 14(4): p. 417–419.

11. Soneson, C., M.I. Love, and M.D. Robinson, Differential analyses for RNA-seq: transcript-level estimates improve gene-level inferences. F1000Res, 2015. 4: p. 1521.

12. Wickham H (2016). ggplot2: Elegant Graphics for Data Analysis. Springer-Verlag New York. ISBN 978-3-319-24277-4, h.g.t.o.

13. Warnes, G.B., B. & Bonebakker, L. & Gentleman, R. & Huber, W. & Liaw, A. & Lumley, T. & Mächler, Martin & Magnusson, Arni & Möller, Steffen. (2008). gplots: Various R programming tools for plotting data.

14. Hayer, A., et al., Engulfed cadherin fingers are polarized junctional structures between collectively migrating endothelial cells. Nat Cell Biol, 2016. 18(12): p. 1311–1323.

15. Sanjana, N.E., O. Shalem, and F. Zhang, Improved vectors and genome-wide libraries for CRISPR screening. Nat Methods, 2014. 11(8): p. 783–784.

16. Nicolle, R., et al., Integrated molecular characterization of chondrosarcoma reveals critical determinants of disease progression. Nat Commun, 2019. 10(1): p. 4622.

17. Frassica, F.J., et al., Dedifferentiated chondrosarcoma. A report of the clinicopathological features and treatment of seventy-eight cases. J Bone Joint Surg Am, 1986. 68(8): p. 1197–205.

18. Morley, S.C., The actin-bundling protein L-plastin supports T-cell motility and activation. Immunol Rev, 2013. 256(1): p. 48–62.

19. Schaffner-Reckinger, E. and R.A.C. Machado, The actin-bundling protein L-plastin-A double-edged sword: Beneficial for the immune response, maleficent in cancer. Int Rev Cell Mol Biol, 2020. 355: p. 109–154.

20. Chen, C., et al., AP4 modulated by the PI3K/AKT pathway promotes prostate cancer proliferation and metastasis of prostate cancer via upregulating L-plastin. Cell Death Dis, 2017. 8(10): p. e3060.

21. Ge, X., et al., Exosomal Transfer of LCP1 Promotes Osteosarcoma Cell Tumorigenesis and Metastasis by Activating the JAK2/STAT3 Signaling Pathway. Mol Ther Nucleic Acids, 2020. 21: p. 900–915.

22. Machado, R.A.C., et al., L-plastin Ser5 phosphorylation is modulated by the PI3K/SGK pathway and promotes breast cancer cell invasiveness. Cell Commun Signal, 2021. 19(1): p. 22.

